# Energy-based and energy-free food-consumption are correlated in captive non-human-primates: A novel dispenser for feeding and behavioral enrichment

**DOI:** 10.1101/803528

**Authors:** Yosef Shohat, Rony Paz, Raviv Pryluk, Aryeh H. Taub

## Abstract

Non-human primates (NHP) provide an important model for studying biological mechanisms that underlie behavior and cognition, and are crucial for supplying translational knowledge that can aid the development of new clinical approaches. At the same time, the importance of the 3Rs to minimize suffering during experiments encouraged the development of environmental enrichment programs. Among them, tools for feeding and foraging are central. However, it remains unclear whether the behavioral enrichment tools are used by the animals only for feeding and to satisfy hunger (and hence for survival), or whether these feeding tools serve also as behavioral enrichment in itself (namely, the animals enjoy it per-se). To answer this, we designed a novel dispenser method – that requires significant yet reasonable energetic effort to obtain food - and tested food consumption via the dispenser compared to free-access, namely that did not require any effort on the animal side. We found that primates consumed food from both the dispenser and when presented in free-access, and importantly, that the consumption via the dispenser was in correlation with the consumption in free-access. This was similar across different subjects, different times during the day, and different types of food. We suggest that monkeys can benefit from using the dispenser for food consumption, but also benefit from it for play (i.e. as behavioral enrichment in itself). Such an approach allows non-human-primates to preserve their natural food procurement activities.

## Introduction

The use of nonhuman primates (NHP) in biomedical research is increasing reaching about 100,000 animals a year [1]. NHPs are mainly used to study HIV/AIDS, brain function, neurodegenerative diseases, Immuology, and addiction [2]. In particular, NHPs are a valuable model for studying brain mechanisms of anxiety/mood-disorders such as generalized-anxiety and post-traumatic stress disorder (PTSD) [3–9], as the major implicated brain regions such as the amygdala and the prefrontal cortex are extensively evolved in NHPs making it an appropriate model [10–13]. Therefore, a major concern for creating a valid behavioral model is preventing uncontrolled irrelevant factors from interacting with the induction and manipulation of anxiety of the experimental design.

The lack of environmental enrichment was shown to impair brain development and might lead to anxiety-like behaviors (e.g. stereotypic behaviors) in laboratory animals [14–16]. While it was initially assumed that environmental enrichment can compromise the standardization of laboratory experiments, recent results reject this notion and showed that enrichment can contribute to the animals’ welfare without increasing the variability and without damaging reliability of the main experimental results [17]. Moreover, studies have shown that enriched environment prevent abnormal brain development, reduce stereotypic behavior and anxiety-like behavior in laboratory animals. Therefore, it is now commonly accepted that environmental enrichment is beneficial for the animal and in parallel can increase the validity of biomedical and behavioral studies [16, 17].

A central goal of environmental enrichment in captive animals is to motivate the animals to engage and exercise species-typical behaviors [18]. Although feeding and food-related enrichment in NHPs is traditionally studied in respect to dietary enrichment [18], food procurement activities are actually a central component in monkeys’ daily schedule in the Nature. Therefore, food enrichment can be used to preserve NHP’s natural exploration and foraging behaviors.[19]. To this end we designed and fabricated a novel food dispenser that requires the animal to invest cognitive and physical efforts in order to obtain food. This allowed us to tackle a central question in the field: is food-foraging a result of survival need, or is it also a behavioral enrichment?

To address this, we examined food consumption via the novel dispenser, namely, requiring effort and energy investment, and compared it to obtaining food from an open container, namely without any need for investing effort. We found a correlation between the amount of food consumed via both routes, showing a relationship between the amount of food intake and the behavioral foraging. The results suggest that foraging is not used exclusively to satisfy hunger and/or energy need. We conclude that artificially inducing and manipulating foraging behavior can be used as environmental enrichment in captive HNPs and laboratory animals.

## Methods

### Experimental Model and Subject Details

#### 1.1 Animals

Four adult male macaca fascicularis (4–8 kg) were used for the present study. All, experimental procedures and housing in the non-human primate colony in the Weizmann Institute were approved and conducted in accordance with the regulations of the Weizmann Institute Animal Care and Use Committee (IACUC), following NIH regulations and with AAALAC accreditation.

#### 1.2 Food dispenser

The dispenser (figure 1) is a ~59 cm. tube, made of a transparent PVC with a handle attached either to the left or right side. Rotating the handle by 360° releases a precise food portion that slides through the tube to the cage floor. The floor is covered with wooden chips, and the animal therefore has to descend and search for the food, an action that usually takes up to one minute.

**Figure 1:**
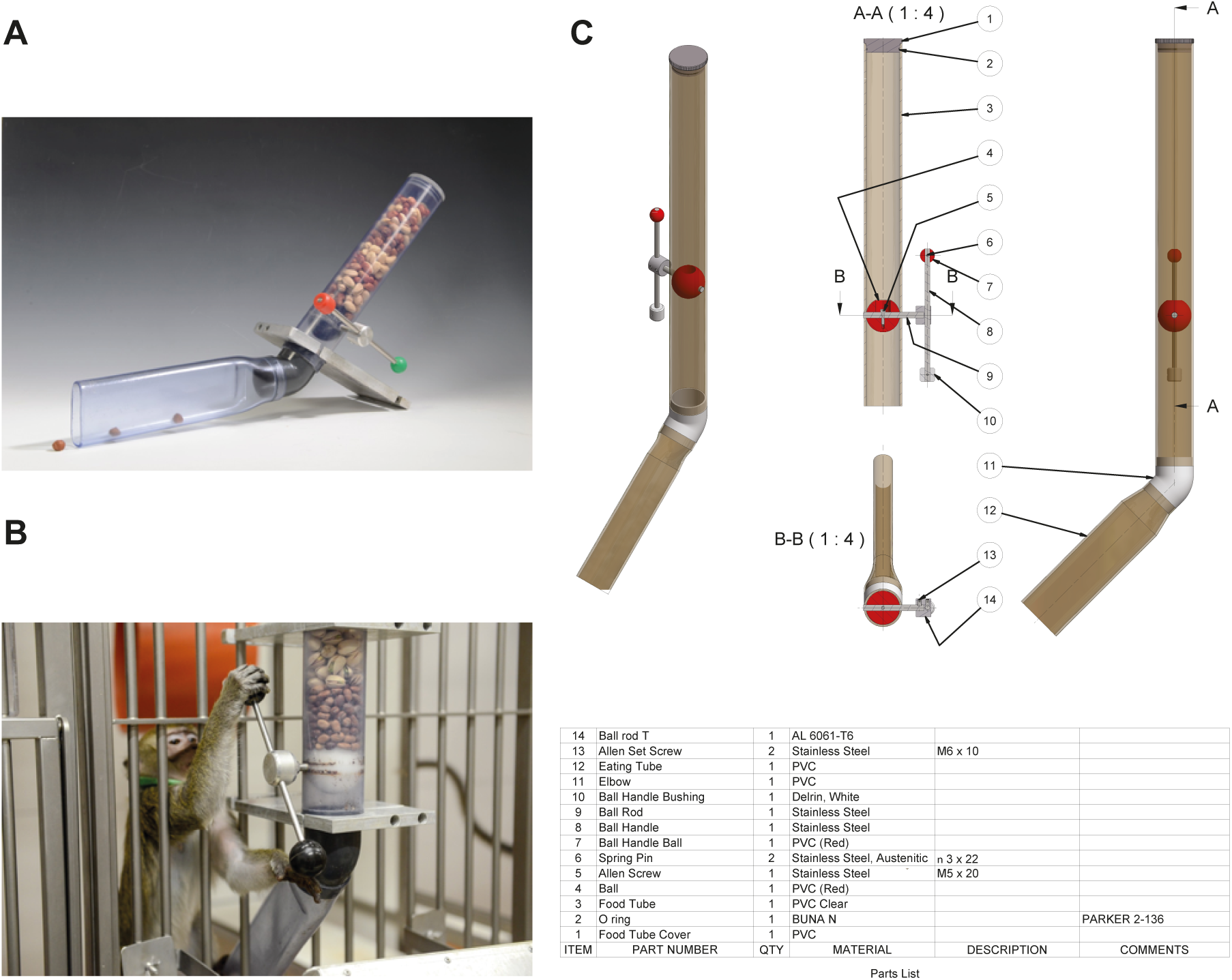
The dispenser. (A – C) A novel dispenser was designed and manufactured (Weizmann workshop) to meet with the physiological characteristics of macaca fascicularis, and can be further adapted to other NHP species. Parts and dimensions are provided in (C).

Key elements in the design of the dispenser were:

1. Using a material (PVC) that ensures the robustness of the instrument.
2. A rotary mechanism with balanced friction that forces the monkey to invest physical effort on one hand, but will allow success of food extraction without inducing frustration on the other.
3. A water proof instrument.
4. A solid fixing mechanism adapted to the animal cage.
5. A transparent tube that allows the monkey to see the type of food.
6. Opening the food intake by the caregiver for re-filling it is convenient and easy.

#### 1.3 Experimental cage

The homecage at the Weizmann facility is 7 × 5 × 2.5 m. (L, W, H), constructed of 6 chambers and divided by removable partition. For the experimental arena we used a chamber of 2 × 1.5 × 2.5 m. (L, W, H).

#### 1.4 Daily session

Two pairs of animals were kept throughout the study.

Each daily session lasted four (4) hours in duration, either from 08:00 – 12:00 or from 14:00 – 18:00. During the four hours of experiment, 200 grams of a single food type, either monkey dumplings, sunflower seeds, or watermelon seeds, were provided either by placing the amount in the cage with free access, or in the dispenser.

The monkeys were familiarized during several weeks (> 20) prior to the beginning of the study. During the day and during the experiment, the dry food was given only in the experiment cage and additional fruit and vegetables were given in a different cage.

On each experimental day, a different food type was provided via the dispenser and free-access (figure 2A, n_food type_ = 14, for a total of 42 days).

**Figure 2:**
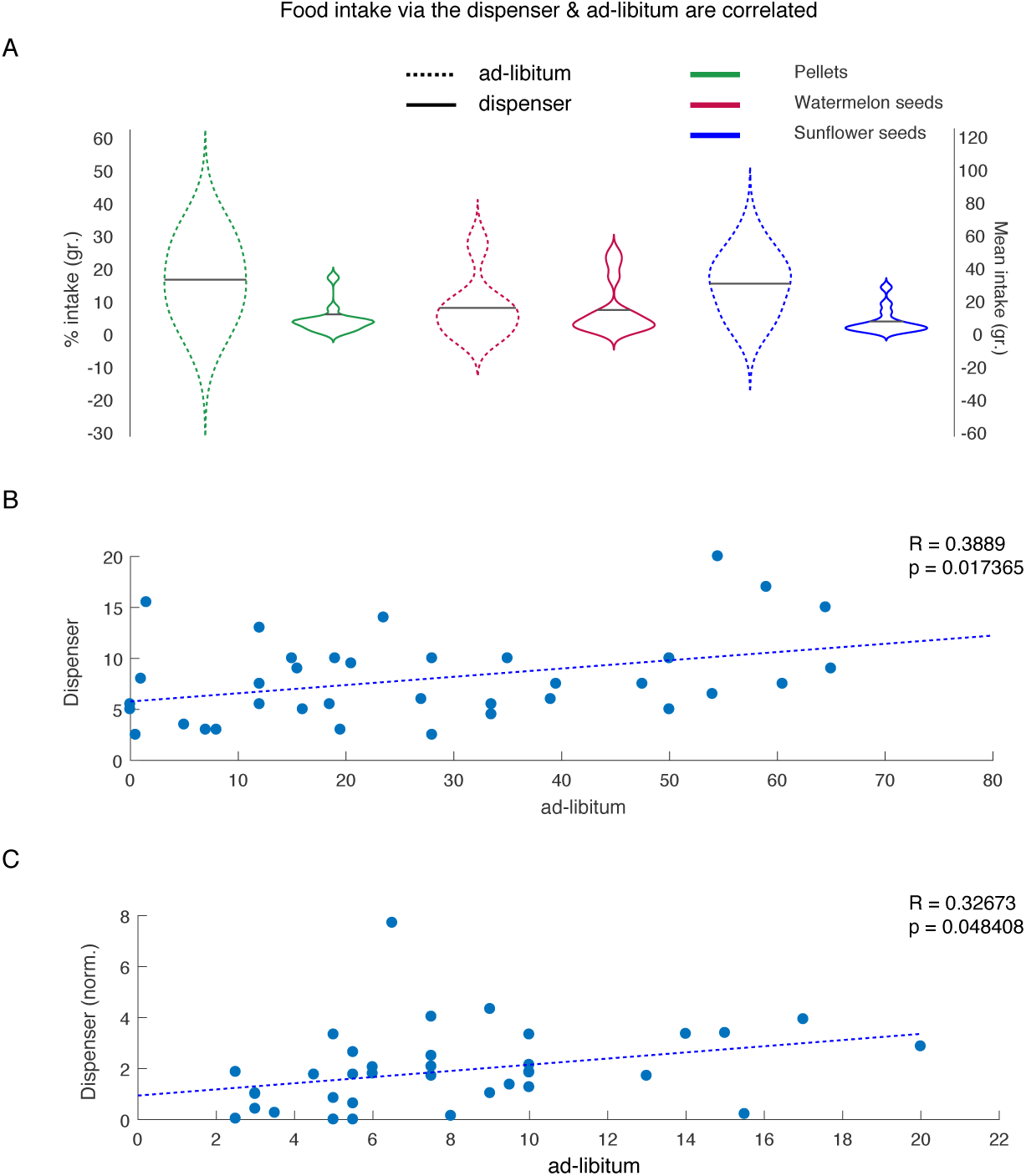
Food intake via the novel dispenser is proportional to that of free-access. (A) mean percent food intake par day and food type (y-axis left) and mean intake of food presented as absolute numbers (y-axis right) both indicate increased food consumption in the free-access route. Black lines indicate mean intake. (B) when examining the relationship in intake between experimental conditions (presented as mean gr./day), we found a strong correlation between free- access and via the novel dispenser. (C) The correlation remained after normalization of the intake via the dispenser to the amount of food available per lever turn.

#### 1.5 Statistics

##### Normalization of dispenser output

To normalize the dispenser output and control for different food size, to equate the amount of food delivered with each lever turning, we computed the number of turns needed to empty the food tube (averaged over 10 times for each food type).

## Results

In order to compare the intake of food between free-access and the dispenser, we computed the mean daily consumption over days for each type of food (figure 2). Results showed increased intake in free-access for pellets (34.5 ± 1.2 gr. SEM, 17.3% of the offered 200gr) vs. dispenser (13.4 ± 1.1 gr., 6.7%), watermelon seeds 17.3 ± 1.4 gr. (8.6%) vs. 16.1 ± 1.2 gr. (8%), sunflower seeds 32.17 ± 0.7 gr. (16.1%) vs. 9.0 ± 0.5 gr. (4.5%). The daily mean consumption over days and food types was significantly higher for the free-access (28.1 ± 3.2 gr., 14.1%) than for the dispenser (12.9 ± 2.4 gr., 6.4 %) (p = 0.0003, two way anova, main effect for condition, df = 1, f = 14.31; no significant effect of food type, p = 0.33, df = 2, f = 1.1) or of delivery method in the watermelon seeds (p = 0.72, post-hoc hsd test).

We further found strong correlation in daily consumption between the two delivery-methods, namely via the dispenser and free-access (figure 2B; r = 0.388, p = 0.01), as well as for the normalized mean consumption between the two delivery methods (figure 2C; r = 0.326, p = 0.04, both Pearson correlation). Taken together these results indicate that the amount of food via both methods was proportional and suggests that although monkeys had free-access to food, they still used the dispenser to reach full satiation.

In order to control for the possibility that these effects are due to differential food consumption by only one of the monkey pairs in the study, we compared the mean consumption of food by free-access vs. dispenser between monkey pairs (figure 3A, 70.7 ± 4.79% and 29.29 ± 2.78% vs. 65.77 ± 4.16 % and 34.23 ± 4.1% SEM, for free-access vs. dispenser, respectively). As expected, one-way anova analysis of daily food intake by monkey pair was not significant (percent intake: p = 0.42, df = 1, f = 0.66, actual intake in grams: p = 0.15, df = 1, f = 2.2), whereas differences between free-access and dispenser were significant (p = 0.0025, df = 1, f = 5.2). We further controlled for the possibility that differences can be attributed to the time-of-day in which the measurements were taken (figure 3B, percent intake: p = 0.12, df = 1, f = 2.41, actual intake in grams: p = 0.3, df = 1, f = 1.08). Here again, differences between free-access and dispenser were significant (p < 0.0001, df = 1, f = 10.21).

**Figure 3:**
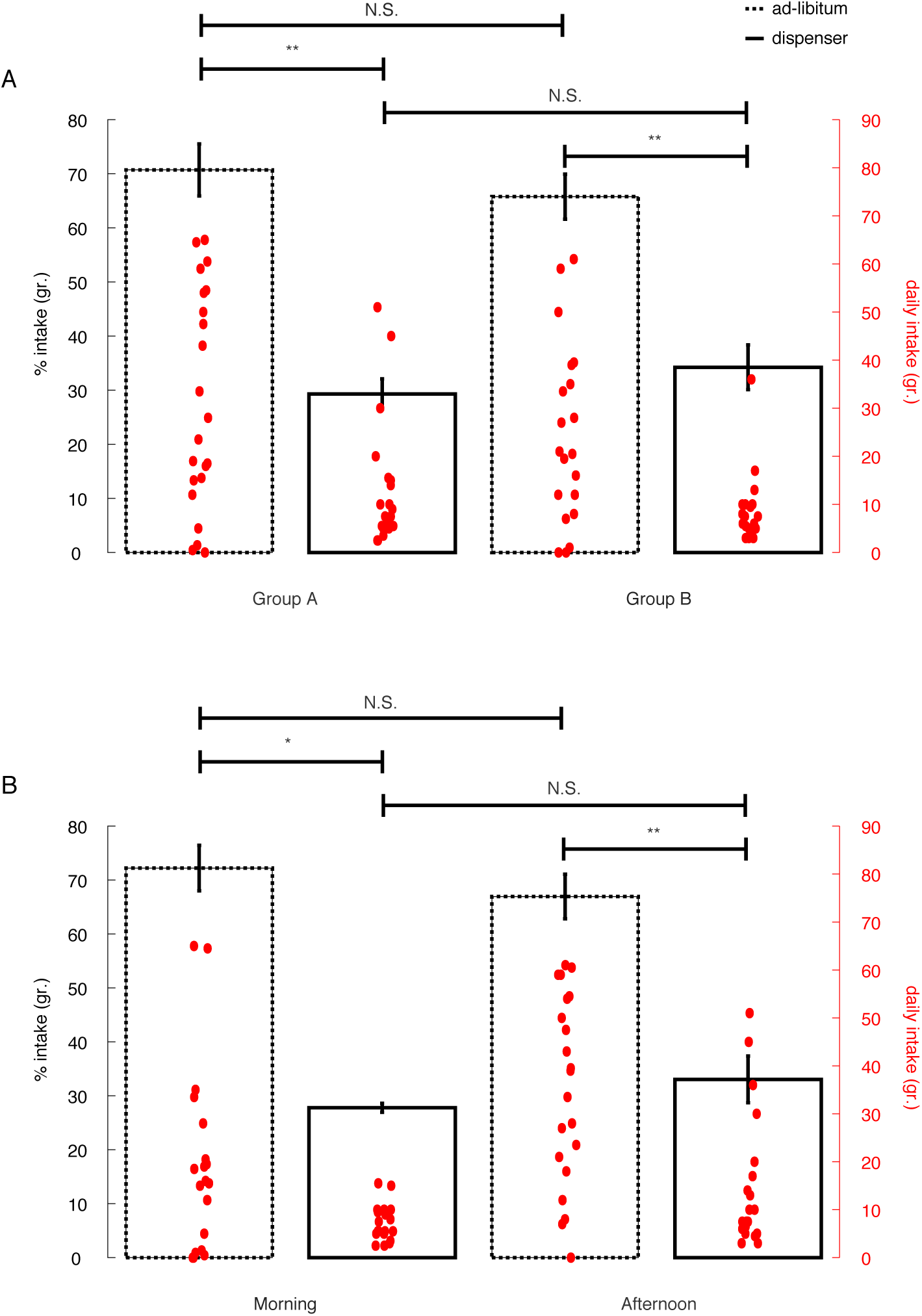
Food intake is similar across animal groups and daily schedule. (A) Food intake for both experimental conditions, measured both as mean percent in grams and absolute intake (grs.) was similar across pairs of animals used in the present study. Whereas no significant differences were found between NHPs groups for free-access (p = 0.6) and dispenser (p = 0.9), differences between ad-libitum and dispenser were significant within each group (group A: p = 0.008, Group B: p = 0.01). (B) Food intake for both experimental conditions was not affected by the time-in-day. Whereas no differences were found for morning vs. afternoon for free-access (p = 0.18) and dispenser (p = 0.06), differences were significant between free-access and dispenser for within each experimental time-of-day (morning: p = 0.03, afternoon: p = 0.007). Data presented as mean ± SEM. * p < 0.05; ** p < 0.01.

## Discussion

This study was aimed at answering a central question in the field of animals’ enrichment: Do primates devote time and energy to acquire food, even if the same food is available without effort? Our findings strongly suggest that although more food was consumed when effort was not required, the monkeys nevertheless invested energy and time to get significant proportion of their food through a dispenser that requires energy investment. This result was robust across different types of food, the time during the day (morning vs. afternoon), and individual animals. Altogether, the results suggest that monkeys in captivity are willing, or even prefer, a certain degree of challenge in their search for food.

This idea is supported by the positive and significant correlation between food consumption via the dispenser and the free-access, strongly suggesting that the monkeys used the food they consumed via the dispenser to satiate hunger. In other words, it is highly unlikely that the monkeys used the dispenser primarily for play rather for food consumption. An additional observation was that across sessions and days, a significant amount of food still remained available by the time session has ended, strongly suggesting that the dispenser was not used due to lack or low availability of free-access food. Taken together, we suggest that the dispenser was used not only as a toy or play, but also for nutrition and to satiate hunger.

Since the amendment to the Animal Welfare Act (1985), considerable effort have been expanded to address the legal and ethical issues in general, and to develop environmental enhancement programs that include feeding tools in particular [17–19]. There are two main reasons for that, first, the welfare of the animals, and second, enabling the experimental conditions and the animal physiological and mental state to be as relevant as possible to humans, promoting translatability.

To preserve and mimic the natural foraging of primates, we designed a novel dispenser. This dispenser requires investing reasonable energy to acquire food, as in Nature. The dispenser is a basic and affordable tool that can be easily fitted to any primate cage. Indeed, our findings show that the monkeys used the dispenser mimicking their natural foraging behavior. We suggest that using such a dispenser among other enrichment tools will improve the animal’s welfare, and increase the validity of experimental results in translational experiments. Future studies should further examine biochemical and physiological aspects of the animals while they use such a dispenser.

## References

1. Chatfield, K. and D. Morton, The Use of Non-human Primates in Research, in Ethics Dumping: Case Studies from North-South Research Collaborations, D. Schroeder, et al., Editors. 2018, Springer International Publishing: Cham. p. 81–90.

2. Mitchell, A.S., et al., Continued need for non-human primate neuroscience research. Current biology: CB, 2018. 28(20): p. R1186–R1187.

3. Klavir, O., R. Genud-Gabai, and R. Paz, Functional connectivity between amygdala and cingulate cortex for adaptive aversive learning. Neuron, 2013. 80(5): p. 1290–300.

4. Livneh, U. and R. Paz, Amygdala-prefrontal synchronization underlies resistance to extinction of aversive memories. Neuron, 2012. 75(1): p. 133–42.

5. Taub, A.H., et al., Oscillations Synchronize Amygdala-to-Prefrontal Primate Circuits during Aversive Learning. Neuron, 2018. 97(2): p. 291–298.e3.

6. O’Neill, P.-K., F. Gore, and C.D. Salzman, Basolateral amygdala circuitry in positive and negative valence. Current Opinion in Neurobiology, 2018. 49: p. 175–183.

7. Pryluk, R., et al., Shared neural codes for eye-gaze and valence. bioRxiv, 2019: p. 736462.

8. Paton, J.J., et al., The primate amygdala represents the positive and negative value of visual stimuli during learning. Nature, 2006. 439(7078): p. 865–870.

9. Taub, A.H., Y. Shohat, and R. Paz, Long time-scales in primate amygdala neurons support aversive learning. Nature Communications, 2018. 9(1): p. 4460.

10. Ghashghaei, H.T., C.C. Hilgetag, and H. Barbas, Sequence of information processing for emotions based on the anatomic dialogue between prefrontal cortex and amygdala. Neuroimage, 2007. 34(3): p. 905–23.

11. Barton, R.A. and J.P. Aggleton, Primate evolution and the amygdala, in The amygdala: a functional analysis, J.P. Aggleton, Editor. 2000, Oxford U. Press.

12. Pryluk, R., et al., A Tradeoff in the Neural Code across Regions and Species. Cell, 2019. 176(3): p. 597–609.e18.

13. Likhtik, E. and R. Paz, Amygdala-prefrontal interactions in (mal)adaptive learning. Trends Neurosci, 2015. 38(3): p. 158–66.

14. van Praag, H., G. Kempermann, and F.H. Gage, Neural consequences of environmental enrichment. Nat Rev Neurosci, 2000. 1(3): p. 191–8.

15. Wurbel, H., Ideal homes? Housing effects on rodent brain and behaviour. Trends Neurosci, 2001. 24(4): p. 207–11.

16. Wolfer, D.P., et al., Cage enrichment and mouse behaviour. Nature, 2004. 432(7019): p. 821–822.

17. Newberry, R.C., Environmental enrichment: Increasing the biological relevance of captive environments. Applied Animal Behaviour Science, 1995. 44(2): p. 229–243.

18. Huber, H.F. and K.P. Lewis, An assessment of gum-based environmental enrichment for captive gummivorous primates. Zoo Biol, 2011. 30(1): p. 71–8.

19. Lutz, C.K. and M.A. Novak, Environmental enrichment for nonhuman primates: theory and application. Ilar j, 2005. 46(2): p. 178–91.

